# Stream-Specific Feedback Inputs to the Primate Primary Visual Cortex

**DOI:** 10.1101/2020.03.04.977264

**Authors:** Frederick Federer, Seminare Ta’afua, Sam Merlin, Mahlega S. Hassanpour, Alessandra Angelucci

**Author notes:** Corresponding author and Lead Contact. Corresponding author’s address: 65 Mario Capecchi Drive, Salt Lake City, UT 84132, USA. Equal first author contribution (Frederick Federer, Seminare Ta’afua).

## Abstract

The sensory neocortex consists of hierarchically-organized areas reciprocally connected via feedforward and feedback circuits. Feedforward connections shape the receptive field properties of neurons in higher areas within parallel streams specialized in processing specific stimulus attributes. Feedback connections, instead, have been implicated in top-down modulations, such as attention, prediction and sensory context. However, their computational role remains unknown, partly because we lack knowledge about rules of feedback connectivity to constrain models of feedback function. For example, it is unknown whether feedback connections maintain stream-specific segregation, or integrate information across parallel streams. Using selective viral-mediated labeling of feedback connections arising from specific cytochrome-oxidase stripes of macaque visual area V2, we find that feedback to the primary visual cortex (V1) is organized into parallel streams resembling the reciprocal feedforward pathways. These results suggest that functionally-specialized V2 feedback channels modulate V1 responses to specific stimulus attributes, an organizational principle that could extend to feedback pathways in other sensory systems.

## INTRODUCTION

In the primate sensory neocortex, information travels along multiple parallel feedforward (FF) pathways through a hierarchy of areas ^1^. Neuronal receptive fields (RFs) in higher-order areas become tuned to increasingly complex stimulus features, with multiple parallel pathways specialized in processing specific attributes of sensory stimuli (e.g. object form versus motion). The role of FF connections in shaping cortical RFs has long been recognized ^2^. In contrast, little is known about the function of feedback (FB) connections, although they have been implicated in top-down modulations of neuronal responses, such as attention ^3,4^, prediction ^5,6^ and sensory context ^7-10^. One limitation to understanding the computational function of FB is that we lack information on the rules of FB connectivity to constrain models of FB function. For example, it is debated whether FB connections are anatomically diffuse and, therefore, unspecific with respect to the functional domains they contact in lower-order areas, or whether they are patterned and functionally specific ^11^. A related question is whether diffuse and unspecific FB connections integrate information across parallel processing streams, or whether these connections maintain stream specificity, like their reciprocal FF pathways. Answering these questions would further our understanding of FB function and allow us to refine and inspire theories of cortical computation, many of which make specific assumptions about the functional specificity, or lack thereof, of FB connections ^12-16^.

Here we have investigated the anatomical and functional organization of FB connections to the primary visual cortex (V1) arising from the secondary visual area (V2) in the macaque monkey. This FB pathway is well suited to address questions of anatomical and functional specificity and parallel FB pathways, because V2 is partitioned into cytochrome-oxidase (CO) stripe compartments that receive segregated FF projections from specific V1 CO compartments and layers ^17,18^, and CO compartments in V1 and V2 have specialized functional properties and maps ^19,20^. It is now well established that CO blobs project predominantly to V2 CO thin stripes, while V1 interblobs project to V2 thick and pale stripes ^21,22^. These projections arise predominantly from V1 layers (L) 2/3, and 4B, and sparsely from 4A and 5/6, with L4B projecting more heavily to thick stripes compared to other stripe types ^23,24^. However, it is debated whether V2-to-V1 FB connections form similar parallel pathways that segregate within V1 CO compartments ^25-27^, therefore maintaining stream specificity, or diffusely project to all compartments ^28^, thus integrating information across parallel streams. This controversy is primarily due to the lack, in previous anatomical studies, of sensitive anterograde neuroanatomical tracers capable of labeling V2-to-V1 axons fully and selectively, without also labeling the reciprocal V1-to-V2 FF projections. Here, using selective viral-mediated labeling of FB connections arising from specific V2 CO stripes, we show that, like V1-to-V2 FF connections, V2 FB connections to V1 are organized into multiple parallel streams that segregate within the CO compartments of V1 and V2. These results suggest functionally specialized FB channels.

## RESULTS

To selectively label FB connections to V1 arising from specific V2 CO stripes, in 3 animals (7 injections) we first identified the V2 stripes *in vivo*, using intrinsic signal optical imaging. We then targeted to a particular V2 stripe type injections of a mixture of Cre-expressing and Cre-dependent adeno-associated virus, serotype 9 (AAV9), carrying the gene for green fluorescent protein (GFP) or tdTomato (tdT) (see Methods). In 2 animals (6 injections), injections were made blindly with respect to V2 stripe identity, and the stripe location of the injection sites determined postmortem. We have previously shown that in the primate visual cortex, this viral vector combination results in selective anterograde infection of neurons at the injected V2 site and virtually no retrograde infection of neurons in V1 ^8^. We quantified the resulting distribution of labeled FB axons across V1 layers and CO compartments. We present results from a total of 13 viral injections made in area V2 of 5 macaque monkeys, spanning all V2 layers. First, we describe the distribution of V2 FB axon terminals across V1 layers (n=4 cases), and subsequently their distribution relative to CO blobs and interblobs (n=9 cases).

### Laminar Distribution of V2 Feedback Projections to V1

Here we present results from 4 injections of the anterograde viral mixture that were made blindly into V2. Alignment of the injection sites to V2 sections stained for CO (see Methods) revealed that two AAV9-tdT injections were confined to thin stripes (one example is shown in **Fig. 1A-B**) and two AAV9-GFP injections encompassed all 3 stripe types, but were mostly confined to pale-lateral stripes (one example is shown in **Fig. 1A,C**). Following Federer et al. ^23^, here we term pale stripes located lateral or medial to an adjacent thick stripe as pale-lateral or pale-medial, respectively. Analysis of the laminar distribution of labeled FB axons resulting from these injections was performed on V1 tissue sectioned perpendicular to the layers along an axis parallel to the V1/V2 border. Fluorescent label was first imaged in sample sections, the same sections were then stained for CO, to reveal layers, and finally the sections were re-imaged simultaneously for both CO and fluorescent signals, to maintain perfect alignment of the FB label and CO-defined cortical layers (see Methods). Qualitative observations of sections imaged for GFP and tdT fluorescence (**Fig. 1D** and left panels of **Fig. 1E-F**), and quantitative analysis of fluorescent signal intensity across layers (right panels in **Fig. 1E-F**; see Methods for analysis details) demonstrated that V2 FB neurons terminate most densely in L1 and the upper part of L2, and in L5/6 (particularly 5B and 6). There were also weaker FB projections to the lower part of L2 and upper part of L3. A sparser projection to L4B, instead, was seen only after injections that encompassed thick and pale stripes, but not after injections confined to thin stripes. Little or no projections were seen in L4A and 4C. In the layers devoid of terminal FB axons, nevertheless, the axon trunks of labeled FB axons could be seen ascending vertically to the upper layers, but these axons did not send lateral branches in these layers.

**Figure 1.**
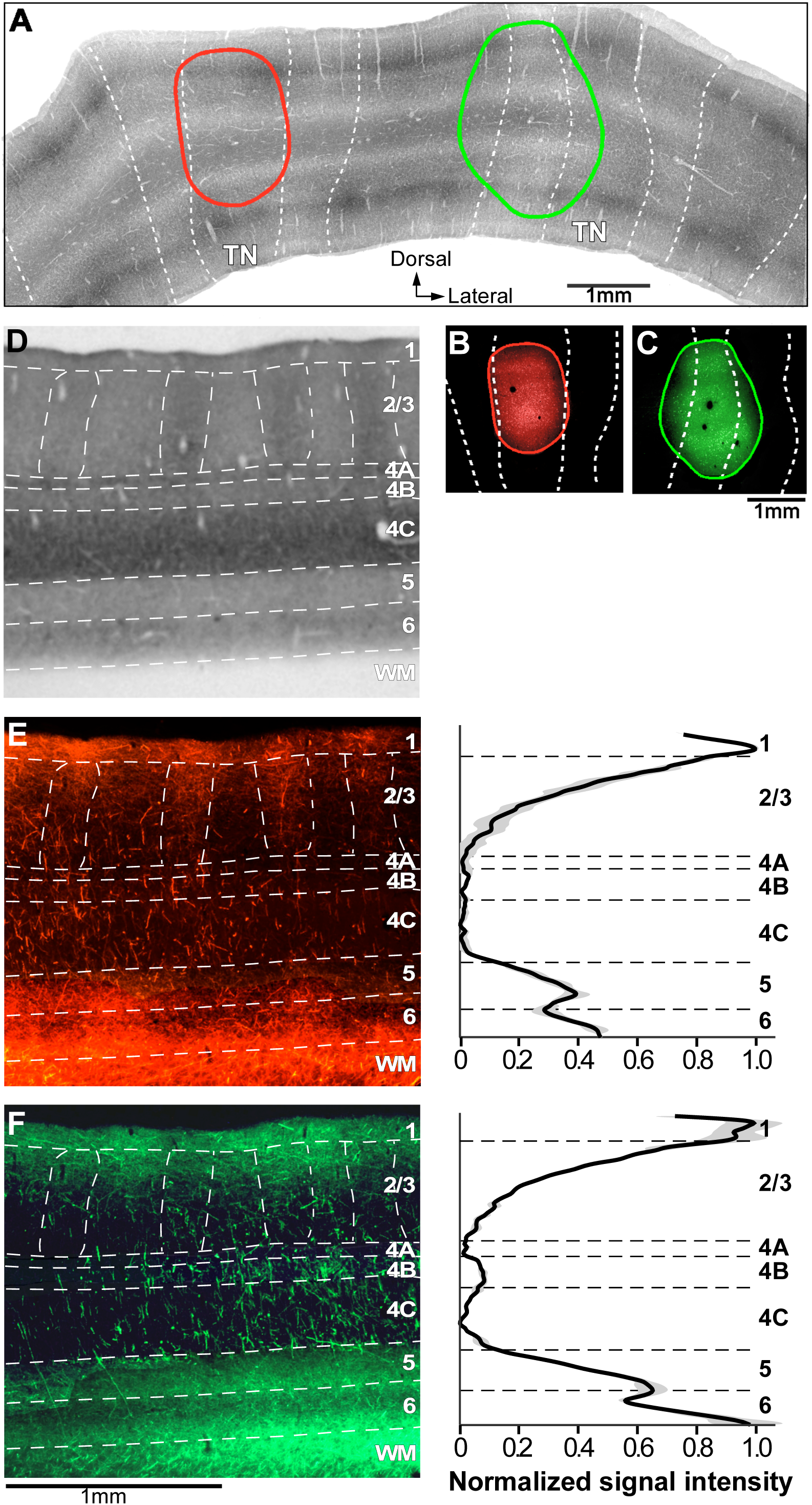
Laminar Distribution of V2 Feedback Projections to V1. Case MK359LH. **(A)** Micrograph of a merged stack of V2 sections stained for CO, to reveal the stripes (delineated by *dashed white contours*). *Red and green ovals* are the composite outlines of an AAV9-tdT and AAV9-GFP injection site, respectively, taken from (B-C) and overlaid onto the V2 CO stack. *TN:* thin stripes. **(B-C)** Micrographs of an AAV9-tdT and an AAV9-GFP injection site, respectively, taken under fluorescent illumination. While the images are from a single tissue section, the *red and green outlines* mark the full extent of the injection sites through the layers, and were generated by aligning images of injection sites through the depth of the cortex. **(D)** Image of CO-stained section through the layers of V1 (indicated on the right side). *White dashed* contours delineate layer boundaries, and the CO blobs in L2/3. The same tissue section, but imaged for fluorescent label, is illustrated in (E) and (F). **(E)** <u>LEFT:</u> Image of tdT-labeled FB axons through V1 layers resulted from the thin stripe injection shown in panels (A-B). <u>RIGHT</u>: Population average of normalized fluorescent signal intensity across V1 layers following thin stripe injections (n=2). **(F)** Same as (E) but for FB label resulted from injections (n=2) that were centered on a pale-lateral stripe, but spilled into adjacent thick and thin stripes. Scale bar under left panel in (F) is valid for (D) and left panel in (E).

Layer-by-layer statistical comparisons across stripe types (thin vs. all stripes) revealed a significant difference in the projections to L4B, which were virtually absent after thin stripe injections, but present when the injection encompassed also the pale/thick stripes. Moreover, injections in thin stripes resulted in significantly less projections to L6, compared to injections that additionally encompassed thick/pale stripes (p <0.001 for both L4B and L6 comparisons, Mann-Whitney U test). Within each stripe group, we also statistically compared the resulting FB label intensity across the sublayers of L4 (4A, 4B, 4C). We found no significant difference in the amount of FB projections from thin stripes to the different L4 sublayers (**Fig. 1E**). In contrast, following injections that additionally involved pale/thick stripes, L4B showed significantly denser FB terminations compared to L4A and 4C (p < 0.001 and p < 0.003, respectively, Kruskal-Wallis test with Bonferroni correction), while L4A did not differ significantly from L4C. In summary, L4B receives significant FB projections from V2 thick/pale stripes but not from V2 thin stripes, and the latter project less heavily to L6 compared to thick/pale stripes.

### Segregated V2-to-V1 Feedback Pathways

We next asked how FB projections from different V2 stripe types are distributed across the tangential plane of V1. **Figure 2A-C** shows one example case in which, using intrinsic signal optical imaging to reveal functional V2 stripes *in vivo* (see Methods, *Optical Imaging*), an injection of AAV9-GFP was targeted to a thin V2 stripe, and an injection of AAV9-tdT to the adjacent pale-medial stripe. **Figure 2A** shows the orientation difference map in two different, but overlapping, regions of interest (ROI1 and ROI2), and the spatial frequency (SF) difference map (low SF-high SF) for ROI2. In the orientation difference map, thin stripes can be identified as regions having weaker or no orientation responses, and pale stripes as regions with strong orientation responses neighboring a region with weak or no orientation domains. In the SF difference map, domains responsive to low SF are located in the thin and thick, but not pale stripes. Stripe location of the injection sites was confirmed postmortem on CO-stained V2 sections (**Fig. 2B**), which were aligned to the optical maps as described in the Methods and in **Supplementary Fig. 1. Figure 2C** demonstrates good correspondence of V2 stripes as defined in the orientation map, SF map and CO staining. GFP- and tdT-labeled FB projections arising from each of these two injection sites formed terminal patches in the superficial layers of V1 (**Fig. 2D)**. When the GFP and tdT images were merged, it was clear that, in the regions where both FB fields overlapped, GFP- and tdT-labeled patches were interleaved (**Fig. 2D** right panel). Importantly, only axon fibers and boutons, but not somata, were labeled in the FB projection zones, indicating these patches were, indeed, formed by axonal terminations of anterogradely-labeled FB neurons (**Fig. 2E**), but not retrogradely-labeled V1 cells. These results indicated that V2 FB projections arising from distinct stripe types segregate within V1.

**Figure 2.**
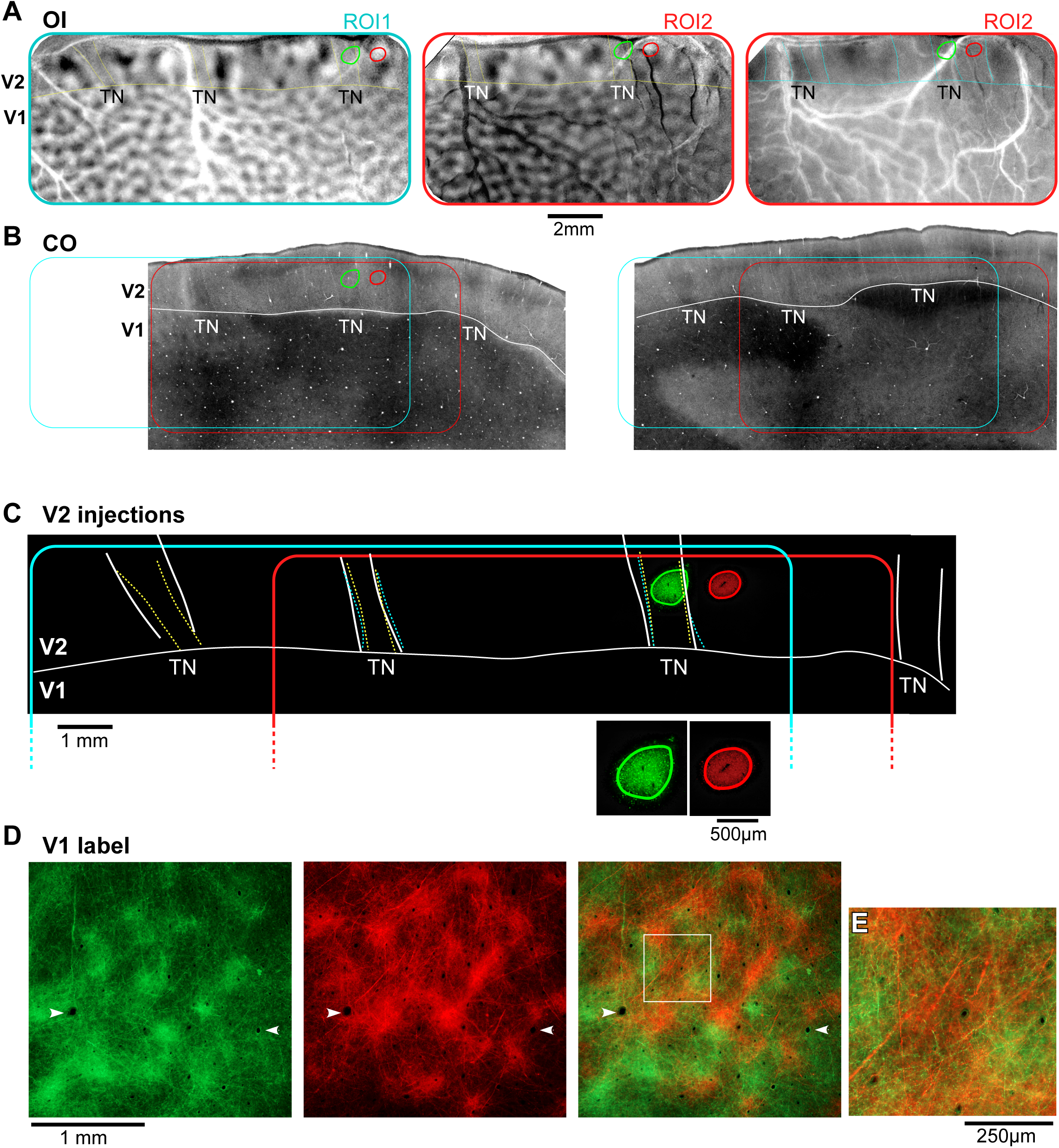
Segregated V2-toV1 Feedback Pathways. Case MK374RHLat. **(A)** Functional maps recorded using *in vivo* intrinsic signal optical imaging (OI). LEFT and MIDDLE: Orientation difference map in two different but overlapping regions of interest (ROI 1 and 2) encompassing V1 and V2. The map in ROI1 was obtained by subtracting responses to achromatic luminance gratings of 135° orientation from gratings of 45° orientation, while the map in ROI2 was obtained by subtracting responses to 0° orientation from those to 90° orientation. *Horizontal solid yellow contour*: V1/V2 border based on the orientation map. *Dashed yellow contours* delineate regions devoid of orientation responses, corresponding to thin stripes (*TN*). RIGHT: SF difference map in ROI2, obtained by subtracting responses to high SF from responses to low SF. *Horizontal solid cyan contour*: V1/V2 border based on the orientation map. *Dashed cyan contours* delineate regions containing low-SF responsive domains (darker regions), corresponding to *TN* and thick stripes. **(B)** Two CO-stained tangential tissue sections, separated by 240 µm (left one is more superficial), encompassing V1 and V2, revealing the stripe pattern and the location of the two injection sites (*ovals*), the latter determined by alignment of fluorescent label to adjacent or same sections stained for CO, using radial blood vessels (see **Supplementary Fig. 1**). *Horizontal solid white contour*: V1/V2 border based on CO staining. *Dashed white contours* delineate *TN* and thick stripes. *Cyan and red box*: locations of optically imaged ROI1 and 2, respectively, on the CO-stained sections. Scale bar in (A) valid for all panels in (A) and (B). **(C)** Micrographs of the AAV9-GFP and AAV9-tdT injection sites, taken under fluorescent illumination, outlined in *green* and *red*, respectively, with superimposed the *TN* stripes outlines from CO (*white*) and OI (*yellow*, orientation, *cyan*, SF). Note good correspondence of CO-defined and functionally-defined stripes. The injection outlines are shown overlaid onto the V2 optical maps in (A) and the V2 CO sections in (B). The injections are shown at higher magnification at the bottom of (C). **(D)** Micrograph of a V1 section through L1B of V1 seen under GFP fluorescence (LEFT), under tdT fluorescence (MIDDLE), and with the two channels merged (RIGHT). *Arrowheads* point to same blood vessels in all panels. GFP and tdT-labeled patches are interleaved. Label inside the *white box* is shown at higher power in (E). **(E)** High power view of FB terminations demonstrating lack of labeled somata, indicating the viral vectors acted as selective anterograde tracers.

We next asked how the terminal patches formed by FB projections arising from each of the stripe types relate to the CO blobs and interblobs of V1, and whether patchy terminations occur throughout the V1 layers of FB termination. Below we present results from a total of 9 viral injections that were confined to V2 thick (n=3), thin (n=3) or pale (n=3) stripes.

#### V2 Thick Stripes Project to V1 Interblobs

Here we present results from a total of 3 viral injections (2 AAV9-GFP, 1 AAV9-tdT, n=3 animals) that were confined to V2 thick stripes. Analysis of the distribution of labeled FB axons relative to the CO compartments of V1 was performed on tissue sectioned parallel to the pial surface. Fluorescent label was imaged in sample sections through the layers where FB axons terminated, namely L1A, 1B, 2/3, 4B and 5/6; the same or adjacent sections were then stained for CO to reveal layers and CO blobs, and imaged. Images of each section were vertically aligned in a sequential stack through the depth of V1, by aligning the radial blood vessels. One example AAV9-GFP injection site confined to a thick stripe is shown in **Figure 3**, and the FB axons in V1 labeled from this injection are shown in **Figure 4 and Supplementary Fig. 2**. The left panel of **Fig. 4A** shows a filtered CO-stained section through V1 L2/3 overlapping the FB termination zone. The CO blob outlines extracted from this image (see Methods) are shown superimposed to the GFP-labeled axons through the depth of V1 (left panels in **Fig. 4B-F**). Qualitative observation of these images revealed patchy FB terminations in V1, with the densest patches located preferentially in the interblobs, across all layers of FB termination. Consistent with the laminar analysis shown above, the densest terminations were located in L1/upper 2 and 5B-6, and sparser projections in L2/3 and 4B. To quantify the distribution of FB projections in the blobs and interblobs of V1, for each layer we measured fluorescent signal intensity within a square region of interest (ROI) centered on each blob overlaying the FB termination zone, and encompassing the average blob and interblob diameter in the CO map (see Methods and **Supplementary Fig. 3A-B**). Fluorescent signal intensity was, then, summed across all blob ROIs within a layer, and normalized to the maximum intensity value, to generate a heat map of fluorescent signals for that layer (**Supplementary Fig. 3C-D, Fig. 4B-F** middle panels). The black circle on each layer heat map shows the average blob diameter estimated from the blobs used to compute that layer heat map. In all layers, the highest fluorescence intensity lay outside the average blob border, indicating the densest FB termination lay in the interblobs. The densest FB label, in this case, was located in the upper right quadrant of the heat maps, rather than uniformly filling the entire interblob region all around the blob. Indeed the densest FB label was consistently located to one side of the blobs, in the termination zone, perhaps due to the small injection site being confined to few orientation domains in the functional map of V2 (**Fig. 3A**), or due to the topographical organization of FB connections which precisely interconnect matching retinotopic sites within the V1 and V2 CO compartments. The heat map for L4B showed much lower fluorescent label intensity compared to the other layers, but branching axons were clearly present in this layer (**Fig. 4E**, left panel), and the highest label intensity was still found within the interblobs (**Fig. 4E**, right panel).

**Figure 3.**
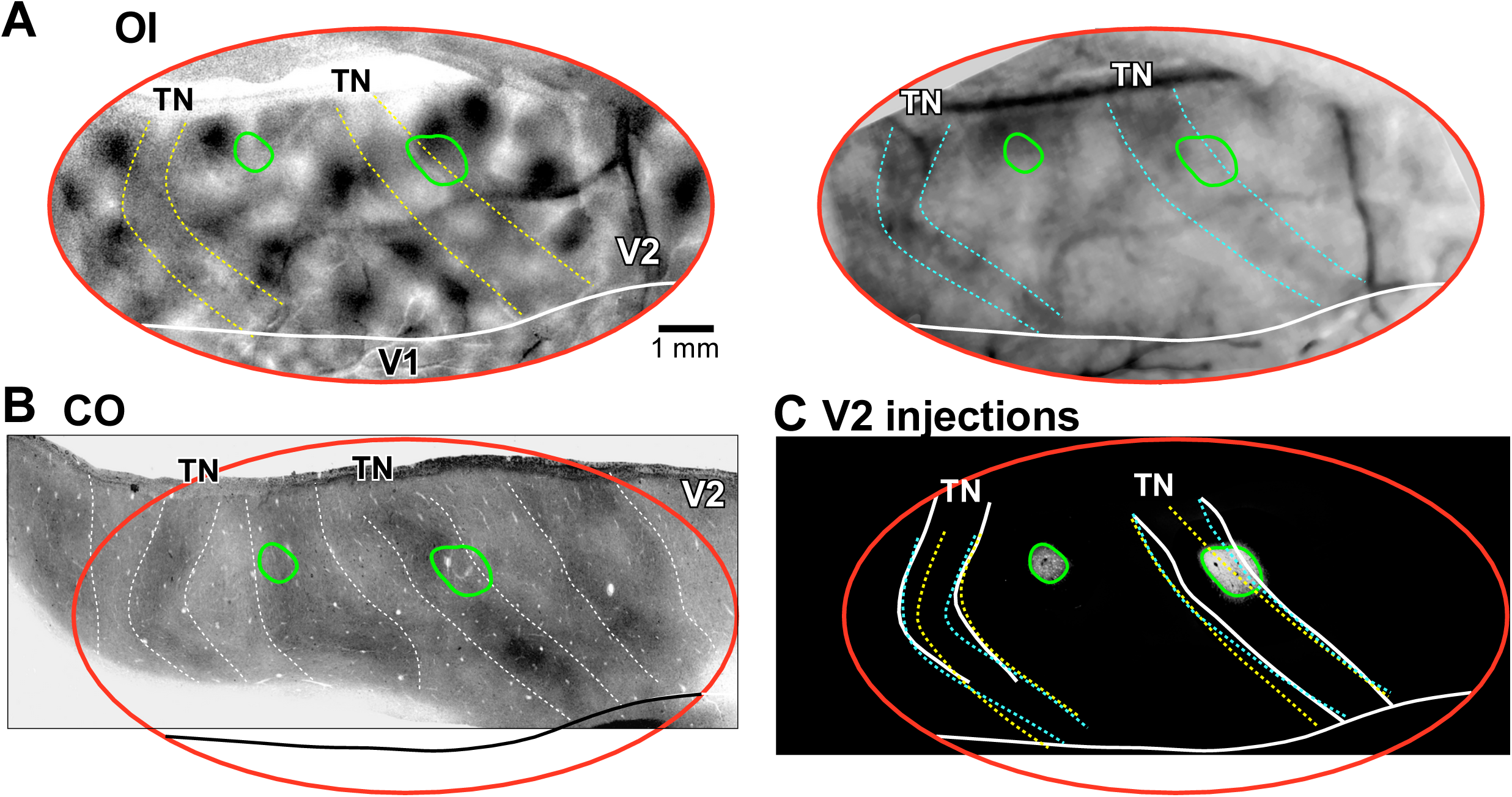
Injection sites in V2 thick and thin stripes. Example of two AAV9-GFP injection sites, one confined to a V2 thick stripe, the other to a thin stripe, in case MK356RH. **(A)** Orientation (45°-135°; LEFT) and SF (low-high; RIGHT) difference maps in V2. *Dashed contours* delineate regions having weak orientation responses (*yellow*), and preferring low SFs (*cyan)*, corresponding to thin stripes; *solid white contour*: V1/V2 border based on the orientation map. **(B)** Stack of aligned and merged tangential V2 tissue sections stained for CO, revealing the stripe pattern. *Red oval*: imaged region from (A), determined by the same alignment procedure described for a different case in **Supplementary Fig. 1**. *Dashed white contours* outline the *TN* and thick stripes; *solid black contour*: V1/V2 border based on the orientation map. **(C)** Micrographs of the two AAV9-GFP injection sites taken under fluorescent illumination and outlined, with superimposed the *TN* stripe outlines from CO (*white*) and OI (*yellow*, orientation, *cyan*, SF). Note good correspondence between functionally-defined and CO-defined stripes. The outlines of the injection sites (*green ovals*) indicate the location of the injection sites on the CO (B) and OI (A) maps.

**Figure 4.**
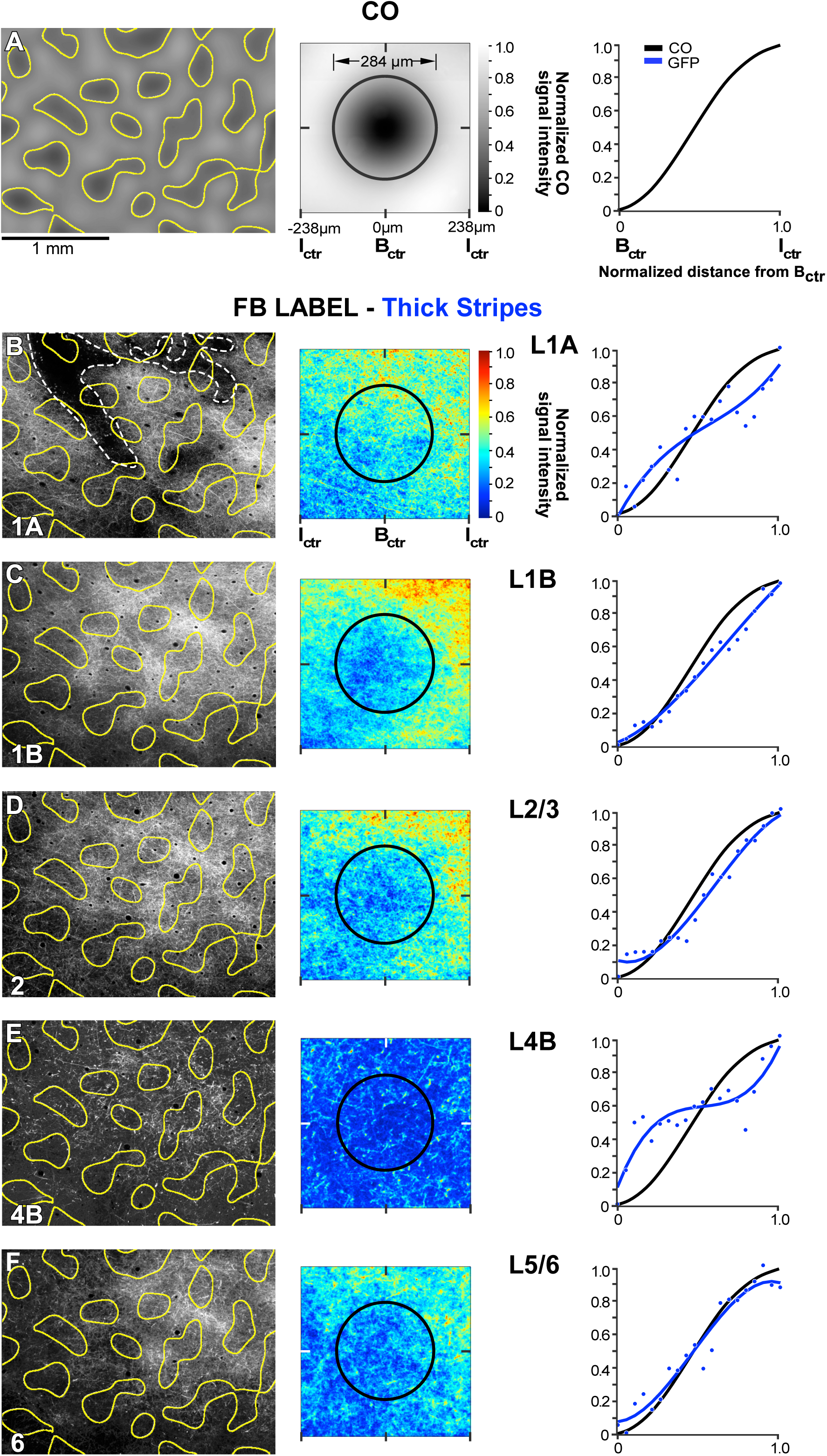
V2 Thick Stripes Project to V1 Interblobs. Case MK356RH. GFP-labeled FB axons in V1 after the thick stripe injection shown in **Figure 3. (A)** <u>LEFT:</u> filtered CO map from a tissue section through V1 L2. Segmenting out the darkest 33% of pixels outlines the CO blobs (*yellow outlines*). Scale bar valid also for left panels in (B-F). <u>MIDDLE:</u> heat map of CO signal intensity, generated by measuring CO intensity within a 476µm^2^ ROI centered around each blob used for the analysis of this case, summing all case ROIs and normalizing to maximum intensity (see Methods and **Supplementary Fig. 3B**). The average blob diameter (*black circle*) for this case was 284µm. <u>RIGHT:</u> plot of the normalized CO signal intensity as a function of distance from the blob center (B_ctr_), computed from the CO heat map, as described in the Methods and **Supplementary Fig. 3B,E**. CO intensity is darkest in the B_ctr_ and brightest in the center of the interblob (I_ctr_). The black CO-intensity curve is repeated on each plot in the right panels of (B-F) for comparison with the FB-label intensity curves. **(B-F)** <u>LEFT</u>: GFP-labeled FB axons in different V1 layers (cortical layer is indicated at the bottom left corner of each panel) with superimposed the blob outlines (*yellow)* from (A, left). *Dashed white lines in (B)* outline the pial vessels. <u>MIDDLE</u>: heat maps of the GFP-label intensity for each respective layer, generated by measuring fluorescent signal intensity within an ROI centered on each blob overlaying the FB label in that layer, summing all ROIs, and normalizing to maximum intensity. The size of the ROI in each layer reflected the average blob and interblob size within the FB termination zone in that layer (see Methods and **Supplementary Fig. 3A,C**). *Black circles*: average blob diameter for the blobs used to compute the heat map for that layer. <u>RIGHT</u>: plots of normalized CO (*black curves*) and GFP-fluorescence (*blue curves*) intensities as a function of distance from the B_ctr_, computed as described in the Star Methods and **Supplementary Fig. 3B, D,E**. Same conventions are used in **Figures 5-7**.

From each layer heat maps, we generated graphs of the normalized fluorescence signal intensity as function of distance from the center of the average blob (blue curves in right panels of **Fig. 4B-F**), as described in the Methods and **Supplementary Fig. 3E**. These curves were compared to similarly computed curves of CO signal intensity as function of distance from the center of the average blob (black curves in right panels of **Fig. 4A-F**, which were computed from the CO heat map shown in the middle panel of **Fig. 4A**). In all layers, the CO and fluorescent signal curves showed similar trends, peaking at the largest distance from the blob center, indicating that the highest fluorescent signals coincided with the brightest regions in CO staining, i.e. the interblobs.

The other two thick stripe injection cases showed similar results to the case shown in **Fig. 4**, albeit due to the smaller size of these injections, the resulting V1 FB label was overall less dense and showed even greater specificity for the interblobs. **Figure 7A** summarizes the thick stripe population data for all layers analyzed.

#### V2 Thin Stripes Project to V1 Blobs

Three viral injections (2 AAV9-GFP, 1 AAV9-tdT, n=3 animals) were confined to V2 thin stripes. One example AAV9-GFP injection site and resulting FB label in V1 are shown in **Figure 2**. A second example AAV9-GFP injection site largely confined to a thin stripe (but encroaching into the adjacent pale-lateral stripe) is shown in **Figure 3**, and the FB axons in V1 labeled from this injection site are shown in **Figure 5 and Supplementary Fig. 2**. Analysis of the distribution of FB labeled axons was performed as for the thick stripe injection case described above. Qualitative observation demonstrated patchy FB terminations, densest in V1 L1/upper 2 and 5B-6, and sparser in L2/3, with little or no projections to L4B (**Fig. 5B-F**, left panels). Both qualitative and quantitative analyses revealed that the densest patches of labeled FB axons lay preferentially inside the blobs, although some sparser label also occurred in the interblobs, likely due to the injection site encroaching into the adjacent pale stripe (**Fig. 5B-F**, middle and right panels). The heat map for L4B showed no significant label (**Fig. 5E**, middle panel), with noisy background label in this layer being equally distributed across blobs and interblobs (**Fig. 5E**, right panel). Qualitative observations of L4B revealed only punctate GFP label with little or no lateral axon branching, suggesting this layer mainly contained unbranched axon trunks traveling vertically (i.e. orthogonal to the plane of imaging) towards the superficial layers. This was in contrast to the branched axonal label seen in this layer after both thick (**Fig. 4E**, left panel) and pale (**Fig. 6E**, left panel) stripe injections.

**Figure 5.**
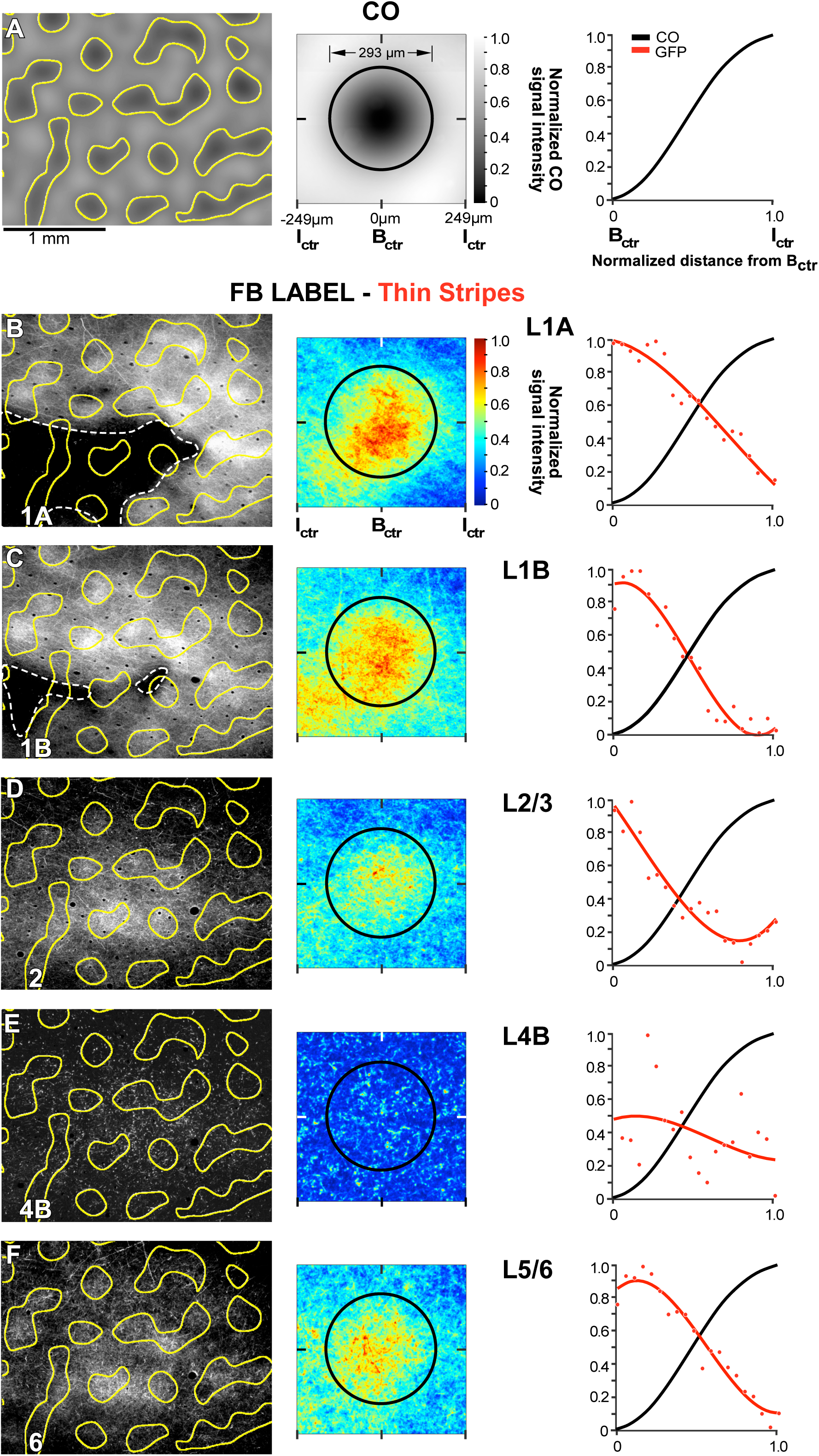
V2 Thin Stripes Project to V1 Blobs. Case MK356RH. Same as in **Figure 4**, but for GFP-labeled FB axons in V1 resulting from the AAV9-GFP injection site confined to a V2 thin stripe shown in **Figure 3**.

**Figure 6.**
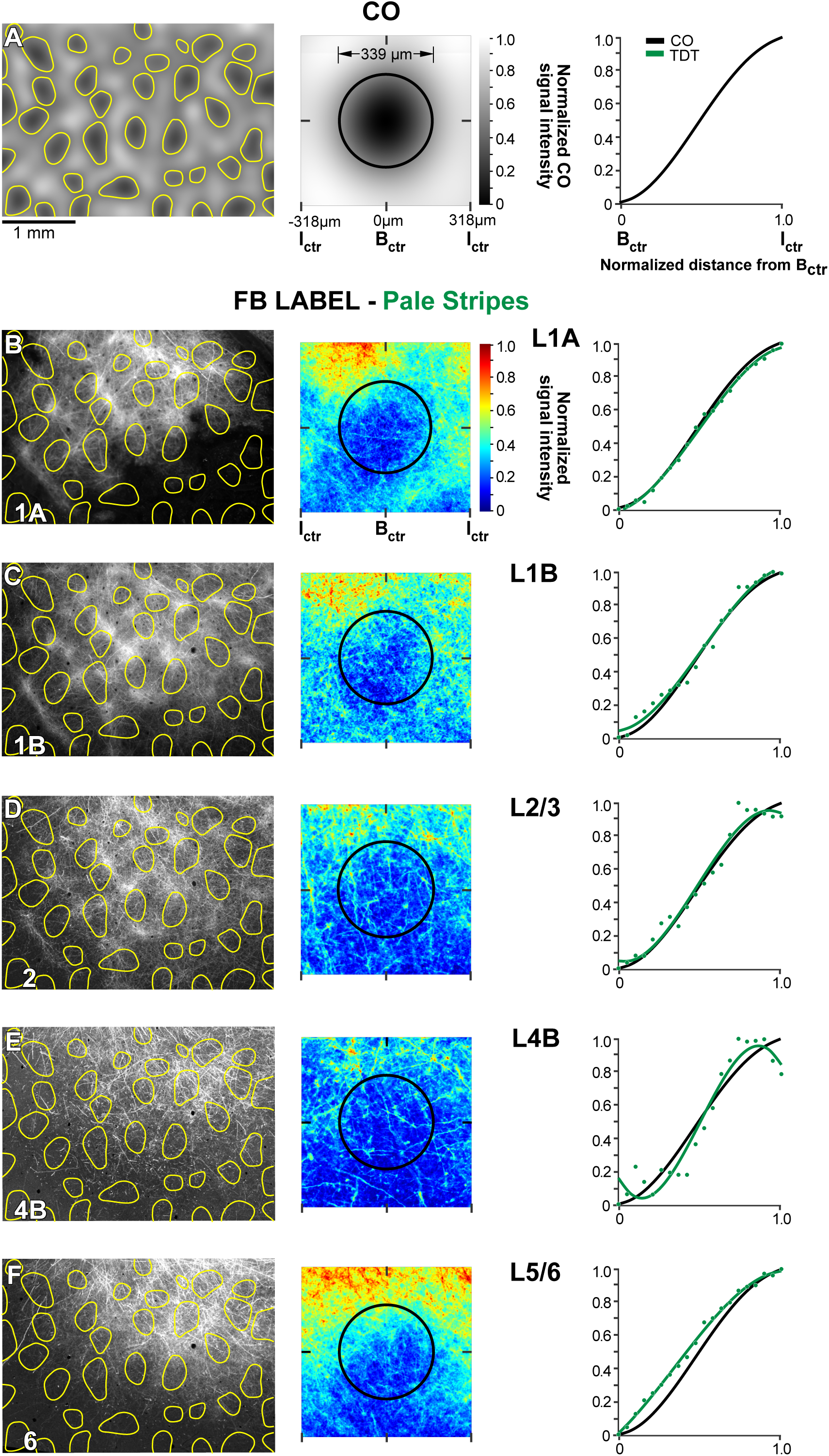
V2 Pale Stripes Project to V1 Interblobs. Case MK374RHLat. Same as in **Figure 4**, but for tdT-labeled FB axons in V1 resulting from the AAV9-tdT injection site confined to a pale-medial stripe shown in **Figure 2A-C**.

The other two thin stripe injection cases showed similar results to the case shown in **Fig. 5**, but since these injection sites were more clearly confined to thin stripes, the terminal FB label showed even greater specificity for blobs. **Figure 7B** shows the thin stripe population data for all layers analyzed.

**Figure 7.**
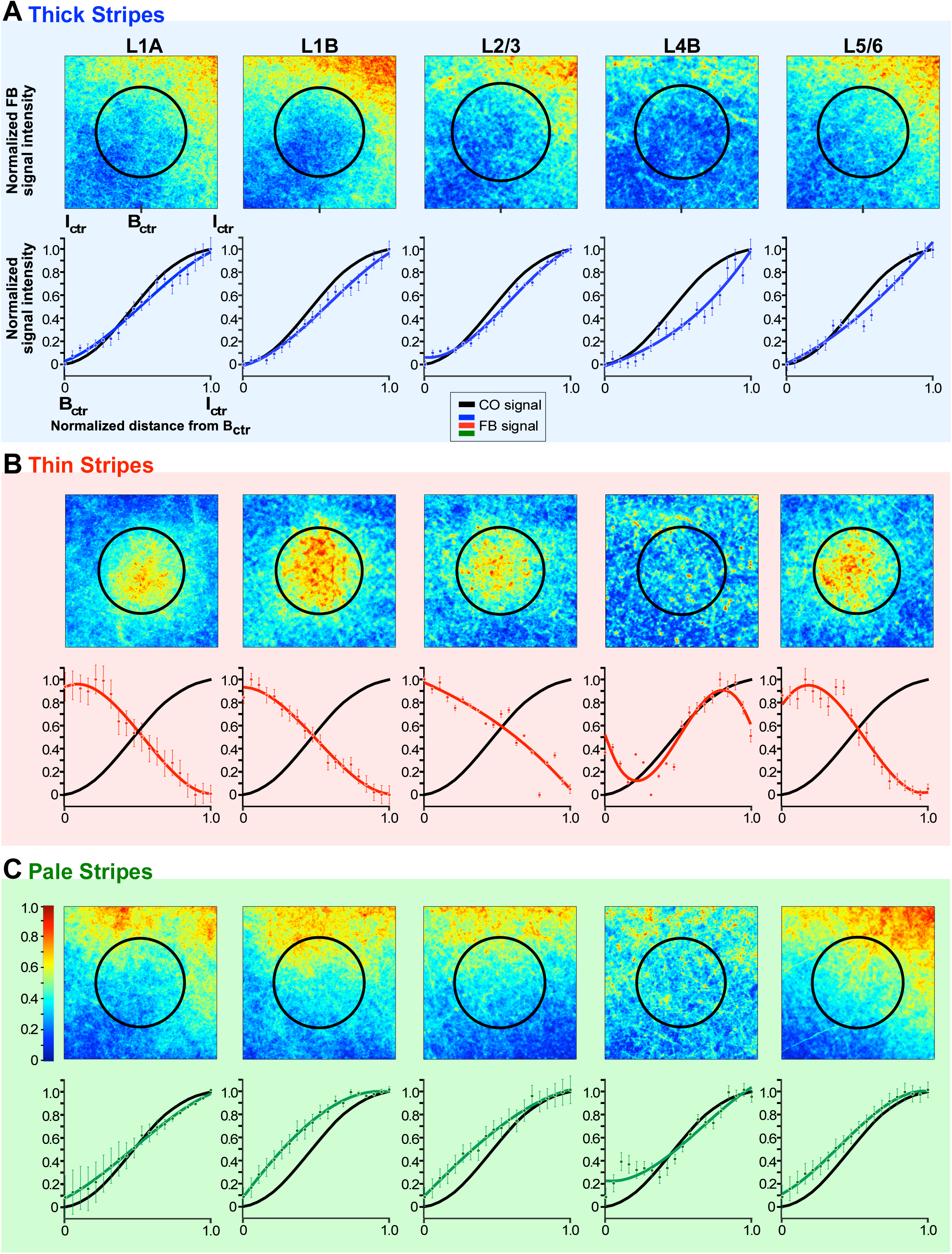
Population Analysis. Heat maps and plots of fluorescent signal intensity, and plots of CO signal intensity for the population of FB projections arising from **(A)** V2 thick stripes (n=3), **(B)** thin stripes (n=3), and **(C)** pale stripes (n=3). For details of how the population averages were computed see Methods. Error bars in the fluorescent signal intensity plots are s.e.m. Conventions are as in **Fig. 4**.

#### V2 Pale Stripes Project to V1 Interblobs

Three viral injections were confined to V2 pale stripes (1 AAV9-tdT in a pale-medial stripe and two AAV9-GFP in a pale-lateral stripe, n=2 animals). One example AAV9-tdT injection site confined to a pale-medial stripe is shown in **Figure 2A-C**, and the FB axons in V1 labeled from this injection site are shown in **Figures 2D-E and 6**. FB axons from this case also showed patchy FB terminations, densest in V1 L1/upper 2 and 5B-6, and sparser in L2/3 and L4B (**Fig. 6B-F**, left panels). FB terminals in L4B formed clear lateral branches indicative of terminations within the layer. Both qualitative and quantitative analyses revealed that the densest patches of labeled FB axons in all layers lay preferentially in the interblobs (**Fig. 6B-F**, middle and right panels). Other pale-stripe injection cases showed similar results. **Figure 7C** summarizes the pale stripe population data for all layers analyzed.

#### Population Analysis

**Figure 7** shows the summary data (see Methods) for the population of injections, grouped by stripe type (n=3 injections per stripe type). Like the individual example cases, the population data shows that FB projections arising from V2 thick and pale stripes terminate preferentially in the interblob regions of V1, across all layers of FB termination (**Fig. 7 A and C**, respectively). In contrast, FB projections arising from V2 thin stripes terminate preferentially in the CO blobs (**Fig. 7B**). Moreover, thick and pale, but not thin, stripes send FB projections to L4B. Statistical comparison of the CO compartment location of densest FB label across stripe groups, for each V1 layer, indicated no significant difference between pale and thick stripes (L1A: p=0.33; L1B: p=0.38; L2/3: p=0.99; L4B: p=0.56; L5/6: p=0.98; Kruskal-Wallis test with Bonferroni correction). In contrast, the CO compartment location of FB projections to V1 differed significantly between thick and thin stripes, as well as between thin and pale stripes, and this was the case for all layers of FB termination (p <0.001 for all comparisons and all layers but L4B for which p =0.01 for the thick versus thin comparison), except for L4B for the thin versus pale stripe comparison which did not reach statistical significance (p= 0.053), likely due to the low sample size. Moreover, when all layers data were pooled together for each stripe group (see Methods), V2 thick and pale stripes showed no significant difference in the V1 CO compartment location of their projections (p = 0.801), while thick versus thin, and thin versus pale stripes were significantly different from each other (p < 0.001 for all comparisons).

## DISCUSSION

We have shown that FB projections arising from the different CO stripes of V2 do not diffusely and unspecifically contact all V1 regions within their termination zones, but rather form patchy axon terminations which largely segregate in the CO compartments of V1: thin stripes project predominantly to blobs and thick and pale stripes predominantly to interblobs. Moreover, all stripe types send densest projections to layers 1/upper 2 and 5B-6, and sparser projections to lower layer 2 and layer 3. However, only thick and pale, but not thin, stripes project to L4B (**Fig. 8**).

**Figure 8.**
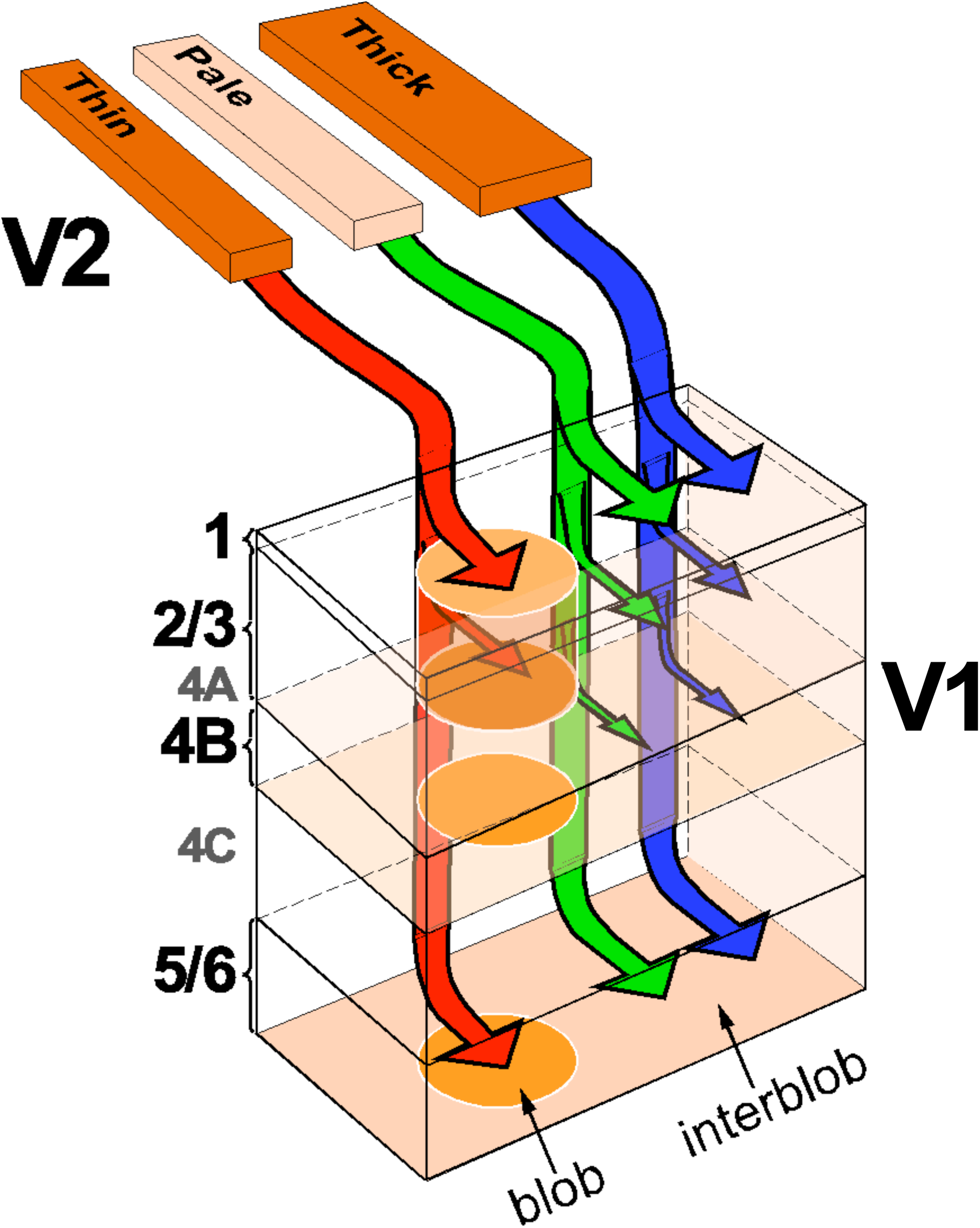
Parallel V2-to-V1 Feedback Pathways. Our proposed model of the FB pathways from V2 to V1, based on the present study. The arrow thickness indicates the relative density of projections to the different V1 layers (L4A is omitted). For all stripe types, layers 1/upper 2 and 5/6 receive the densest FB projections, while lower L2 and L3 receive sparser projections from V2. Both thick (*blue arrows*) and pale (*green arrows*) stripes, but not thin (*red arrows*) stripes send FB projections to V1 L4B. FB projections from thin stripes predominantly target the V1 CO blob columns, while FB projections from thick and pale stripes predominantly target the V1 interblob columns.

There has been a long-standing controversy regarding the clustering and specificity, or lack thereof, of FB connections to V1, mainly attributable to the technical limitations of the neuroanatomical tracers used in previous studies ^11^. Conventional tracers used to label FB axons were either transported anterogradely, but had poor sensitivity and resolution (e.g. tritiated aminoacids, WGA-HRP, and first generation adenoviral vectors), or had good sensitivity, but were transported bidirectionally (CTB, BDA), thus labeling the axons of both FF- and FB-projecting neurons. Studies using the first group of tracers injected in areas V2 or MT concluded that FB connections to V1 are diffuse, i.e. they make non-clustered terminations in their target areas, which are non-specific with respect to the CO compartments or functional domains of V1 ^28-30^. Curiously, however, one such study ^28^ showed at least one example of patchy FB axon terminations in layer 1 of macaque V1, but the authors did not comment on the modularity of FB connections. Studies using injections of bidirectional tracers into V2, instead, provided evidence for clustered and functionally-specific V2 FB projections to V1 ^25,27^. Due to the bidirectional transport of the tracers used in the latter studies, however, it was unclear whether the axon clustering reflected the termination patterns of FB axons or of the axonal collaterals of the reciprocal FF projection neurons, known to form patchy connections ^21,22,31^. For the same reason, it had remained unclear whether FB projections to V1 terminate only in L1 and 6 (which do not send, or only send very sparse FF projections to extrastriate areas), or also terminate in layers that send FF projections to higher cortical areas (such as L2/3 and 4B). In a few studies, FB axons were labeled by bulk injections of PHA-L or BDA in V2 or MT, and reconstructed through serial sections ^32-34^. While these studies were not affected by ambiguity in the interpretation of the origin of axonal label, they, however, provided a limited sample of partially reconstructed FB axons.

To overcome the limitations of previous anatomical studies, in this study we have taken advantage of recent advances in neuroanatomical labeling methods based on the use of viral vectors to deliver genes for fluorescent proteins ^35^. Specifically, we have used a mixture of AAV9-Cre and Cre-dependent-AAV9 to express fluorescent proteins in V2 FB neurons projecting to V1 without simultaneously labeling the axons of the reciprocal V1-to-V2 FF projection neurons. Our results resolve existing controversies by demonstrating unequivocally, in agreement with ^11,27,36^, that FB connections from V2 to V1 form patchy terminations that are predominantly CO-compartment specific. In particular, we demonstrate that in macaque monkey, FB connections arising from the different CO stripes of V2 form parallel channels that mimic the reciprocal parallel pathways from V1 to V2 in this species ^21^. We also show that, with the exception of L1, the layers that give rise to FF connections to V2 (2/3, 4A, 4B, 5/6) receive reciprocal FB inputs. This suggests that V2 FB axons may make direct contacts with V1 neurons sending FF inputs to V2, a hypothesis that we have recently confirmed using rabies virus-mediated monosynaptic input tracing to label monosynaptic inputs to V2-projecting neurons in V1 ^37^. However, while L4B has been shown to send at least some sparse inputs to thin and pale-medial stripes ^23,24,38^, we only observed significant FB inputs to L4B from thick and pale, but not thin, stripes. It is possible that our method of analysis failed to capture very sparsely labeled FB axons in L4B arising from thin stripes. Moreover, our low sample of injections into pale-medial stripes (n=1) did not allow us to assess potential quantitative differences in the FB projections to L4B from the two pale stripe types, as has been previously demonstrated for the L4B-to pale stripe FF projections ^23^. In this study we have not analyzed L4A for potential FB terminations, as this very thin layer is difficult to study in tangential sections, and in pia-to-white matter sections FB projections to L4A appeared to be, at best, very sparse.

Although here we are proposing the existence of at least 3 segregated FB pathways from V2 to V1, our study shows that FB projections from thick and pale stripes both terminate in the interblobs, allowing for the possibility that these projections contact the same V1 cells, therefore integrating information across these two parallel streams. Current studies from our laboratory, using rabies virus-mediated monosynaptic input tracing to label direct inputs to V2-projecting neurons in V1, however, suggest that FB projections from thick and pale stripes contact predominantly distinct V1 projection neurons ^39^.

Segregation of V1-to-V2 FF pathways within CO compartments likely reflects specialized function, because CO compartments in V1 and V2 show specialized neuronal responses properties and functional maps. Specifically, the blob-thin-stripe pathway is thought to process surface properties (color and brightness), the interblob-pale stripe pathway object contours and form, and the interblob-thick stripe pathway object motion and depth ^31,40-51^. Our finding of segregated FB projections within these same CO compartments, therefore, suggests that V2 FB connections to V1 do not integrate information across parallel functional streams, and thus across stimulus attributes, but rather modulate V1 responses to the specific stimulus attributes that are processed within their functional stream, an organization that may support, for example, feature-selective attention ^52,53^, or the feature specificity of extraclassical receptive field effects ^7,54,55^.

Our finding of segregated V2-toV1 FB pathways is consistent with previous results of stream specific organization of corticogeniculate pathways in macaque ^56,57^, suggesting this FB organization may exist at multiple levels of the visual pathway. Our results are also consistent with results in mouse visual cortex, demonstrating segregation of functional FB channels from L5 of higher visual areas AL (anterolateral) and PM (posteromedial) to V1 ^58^, which also matches the functional segregation of the reciprocal FF pathways from V1 to AL and PM ^59^. Modular FB axon terminations have also been demonstrated anatomically in L1 of mouse V1, where FB axons from areas AL and PM form interleaved terminal patches ^60^. The FB patches from AL and AM, in turn align with M2 muscarinic acetylcholine receptor-rich and poor patches, respectively, the latter corresponding to V1 regions specialized for the processing of distinct spatio-temporal features ^60,61^. Interestingly, M2-positive patches are also found in macaque V1 L1 where they are in register with the CO-poor interblobs ^61^. Together with our results, these studies support the notion of functionally specialized parallel FB pathways.

Our results of segregated FB pathways are consistent with predictive coding theories of FB function. According to these theories, the brain generates an internal model of the world, based on sensory data and prior experience, which is refined by incoming sensory data. The latter is compared to the predicted sensory data, and the prediction error ascends up the cortical hierarchy and refines the higher levels’ model ^5,62,63^. Predictive coding models postulate the existence of different neuronal populations encoding expectations and prediction errors, respectively, at each level of the cortical hierarchy. In these models, inter-areal FB connections carry the predictions, while FF connections carry the error in those predictions ^12,13^. In terms of the architecture of hierarchical predictive coding schemes, one key attribute is the functional segregation of conditionally independent expectations, so that descending predictions are limited to prediction errors reporting a particular stimulus attribute, but not prediction errors reporting opposite or different attributes. For example, predictions about the orientation of a visual edge should not project to prediction errors units encoding visual motion, because knowing the orientation of an edge does not, statistically, tell us anything about its motion. This suggests that direct descending predictions will be largely specific for stimulus attributes (e.g., contours vs. motion), which is consistent with our finding of FB-specific channels.

While our results strongly support the existence of functionally-specialized parallel FB pathways, it is noteworthy that our method of analysis was designed to extract the densest FB terminations in V1. However, lighter FB axonal label was visible in the regions between the densest labeled terminal patches. Therefore, we cannot exclude that a smaller population of FB axons shows a different terminal pattern, such as projections to all CO compartments of V1 or projections to the opposite channel. This population of FB axons may serve to integrate information across streams, or to carry FB information about the same stimulus feature to neurons in different V1 CO compartments representing that feature, for example orientation in the blobs and interblobs. Similarly, FB from multiple stripes could affect the same neurons within the same stimulus feature, for example both thick and pale stripes have orientation maps, and FB to V1 from these stripes may be combined on the same V1 cells on the basis of orientation preference.

## METHODS

### Animals

Five adult (4 females, 1 male) cynomolgus macaque monkeys (*Macaca fascicularis)* were used in this study. All procedures involving animals were approved by the Institutional animal Care and Use Committees of the University of Utah and conformed to the guidelines set forth by the USDA and NIH.

### Method Details

AAV9 vectors carrying the gene for either GFP or tdT were injected into specific V2 stripes, which in most cases were identified *in vivo* by intrinsic signal optical imaging. In two animals, instead, injections were made blindly with respect to stripe type. The V1 laminar and CO compartment distribution of resulting labeled FB axons was analyzed quantitatively.

#### Surgical Procedures

Animals were pre-anesthetized with ketamine (10-20 mg/kg, i.m.), intubated with an endotracheal tube, placed in a stereotaxic apparatus, and artificially ventilated. Anesthesia was maintained with isofluorane (0.5–2.5%) in 100% oxygen, and end tidal CO_2_, blood oxygenation level, electrocardiogram, and body temperature were monitored continuously. The scalp was incised, a craniotomy was made to expose the lunate sulcus and about 5 mm of cortex posterior to it (encompassing V2 and a small portion of V1). In two animals (MK359, 4 injections, and MK365, 3 injections) small durotomies were made just caudal to the posterior edge of the lunate sulcus, over V2, and viral vectors were injected into V2. In the reminder of cases (3 animals, total of 12 injections), optical imaging was used to identify the V2 stripes prior to the injections. In the latter cases, a large craniotomy and durotomy (15-20 mm mediolaterally, 6-8mm anteroposteriorly) were performed to expose V2 and parts of V1, a clear sterile silicone artificial dura was placed on the cortex, and the craniotomy was filled with a sterile 3% agar solution and sealed with a glass coverslip glued to the skull with Glutures (Abbott Laboratories, Lake Bluff, IL). On completion of surgery, isoflurane was turned off and anesthesia was maintained with sufentanil citrate (5–10 µg/kg/h, i.v.). The pupils were dilated with a short-acting topical mydriatic agent (tropicamide), the corneas protected with gas-permeable contact lenses, the eyes refracted, and optical imaging was started. Once the V2 stripes were functionally identified (1-3 hrs of imaging, see below), the glass coverslip, agar and artificial dura were removed and the viral vectors were injected in specific V2 stripe types, using surface blood vessels as guidance. On completion of the injections, new artificial dura was placed on the cortex, the craniotomy was filled with Gelfoam and sealed with sterile parafilm and dental cement, the skin was sutured and the animal was recovered from anesthesia. Animals survived 3-4 weeks post-injections, and underwent a terminal 3-5 day optical imaging experiment to obtain additional functional maps. At the conclusions of this experiment the animal was sacrificed with Beuthanasia (0.22 ml/kg, i.v.) and perfused transcardially with saline for 2–3 min, followed by 4% paraformaldehyde in 0.1 M phosphate buffer for 20 min. One case was perfused with 2% paraformaldehyde for 6 min and postfixed overnight.

#### Injections of Viral Vectors

A total of 19 viral vector injections were made in 5 macaque monkeys. We excluded from analysis 6 injections, which either did not produce detectable label in V1 (n=2), or were located in an unidentifiable stripe (n=4). The viral vectors consisted of a 1:1 mixture of AAV9.CaMKIIa.4.Cre.SV40 and either AAV9.CAG.Flex.eGFP.WPRE.bGH or AAV9.CAG.Flex.tdTomato.WPRE.bGH (Penn Vector Core, University of Pennsylvania, PA), which were mixed and loaded in the same glass micropipette (tip diameter 35-45 µm), and pressure injected using a picospritzer. For the cases used to investigate the distribution of FB terminals across V1 layers (**Fig. 1**; n=4), we slowly injected (6-15 nl per min) 105-150 nl of the viral mixture at a cortical depth of 1.2mm from the pial surface. After a 5 min pause, the pipette was retracted to a depth of 0.5 mm, and an additional 105-150 nl of the viral mixture were injected at the same speed. The pipette was left in place for an additional 5-10 min before being retracted. These injection parameters yielded injection sites 1.3-1.8 mm in diameter encompassing all cortical layers. For the cases used to investigate the distribution of FB terminals relative to the V1 CO compartments, we used a similar procedure, but we injected smaller volumes in order to keep the injection site confined to a V2 stripe: 15-30 nl were injected at 1.2mm depth, and an additional 15-30 nl at 0.5-0.6 mm depth. Resulting injection sites ranged in diameter between 0.56 and 0.95 mm and encompassed all V2 layers. One case used for the CO compartment analysis (MK356; **Fig. 3**) received a single injection of 105 nl in a thin stripe and one injection of 75 nl in a thick stripe, both at a cortical depth of 1 mm. The resulting injection sites measured 1.26 and 0.65 mm in diameter, respectively, and encompassed all layers.

Each animal received 2-5 injections in dorsal area V2 of one hemisphere. Within the same hemisphere either vectors expressing different fluorophores (GFP or tdT) were injected, or injections of the same vector were spaced at least 10 mm apart, to ensure no overlap of the resulting labeled fields in V1. In one case (MK356), instead, two injections of the GFP-expressing vector were made in a thick and a thin stripe within the same V2 stripe cycle (**Fig. 3**), which resulted in two labeled fields in V1 that slightly overlapped (see **Supplementary Fig. 2**). In this case, we excluded from analysis the region of label overlap.

#### Optical Imaging

Acquisition of intrinsic signals was performed using the Imager 3001 (Optical Imaging Ltd, Israel) under red light illumination (630 nm). To identify the V2 stripes, we imaged V2 during presentation of gratings varying in orientation and SF. Orientation maps were obtained by presenting for 4s full-field, high contrast (100%), pseudorandomized, achromatic drifting square-wave gratings of eight different orientations at 1.0-2.0 cycles/° SF, moving back and forth at 1.5 or 2°/s in directions perpendicular to the grating orientation. Responses to same orientations were averaged across trials, following baseline correction, and difference images were obtained by subtracting the responses to two orthogonally oriented pairs (e.g. **Figs. 2A, 3A**). Imaging for SF (e.g. **Figs. 2A and 3A**), was performed by presenting high contrast achromatic drifting square-wave gratings of 6 different SFs (0.25, 0.5, 1, 1.5, 2, 3 cycles/°) alternating through 8 different orientations every 500ms, and drifting at 2 cycles/s. To create SF difference maps, baseline corrected responses to high SF (3 cycles/°) were subtracted from baseline corrected responses to low SF (0.25 cycles/°). Baseline correction was performed in 3 different ways and the approach that provided the best maps was selected for analysis: 1) the baseline (pre-stimulus) was subtracted from the single condition response (i.e. the images recorded during stimulation of one stimulus orientation or one SF); 2) the single condition response was divided by the baseline; 3) the single condition response was divided by the “cocktail blank” (i.e. the average of responses to all oriented stimuli or all SF stimuli). Thick stripes were identified as the middle of regions having an orientation-preference map and domains responsive to low SFs; pale stripes as regions having an orientation-preference map neighboring a region with weak or no systematic orientation maps; and thin stripes as regions containing domains responsive to low SF, having weak or no systematic orientation maps (**Figs. 2A, 3A**). In each case, reference images of the surface vasculature were taken under green light (546 nm) illumination, and used *in vivo* as reference to position pipettes for viral vector injections, as well as postmortem to align the functionally identified V2 stripes to the histological sections containing the injection sites and sections labeled for CO to reveal the stripes (e.g. **Figs 2C, 3C and Supplementary Fig. 1**).

#### Histology

Areas V1 and V2 were dissected away from the rest of the visual cortex. The block was postfixed for 1-2 h, sunk in 30% sucrose for cryoprotection, and frozen-sectioned at 40 µm. For the cases used to investigate the distribution of FB terminals across V1 layers (the cases in **Fig. 1**; n=4), the block was cut perpendicularly to the layers and parallel to the V1-V2 border. This sectioning plane allows to easily identify layers, as well as the sequence of V2 stripes, which run perpendicular to the border. For the reminder of the cases (n=9), which were used for the analysis of FB terminals relative to the V1 CO compartments, instead, the block was postfixed between glass slides for 1-2 h to slightly flatten the cortex in the optically imaged area, cryoprotected, and frozen-sectioned tangentially, parallel to the plane of the imaging camera. Sections were wet-mounted and imaged for fluorescent GFP and tdT-labeled cell bodies in V2 (the injection sites) and FB axons in V1 using either Zeiss Axioskop 2 or Axio Imager Z2 microscopes (10x magnification). After digitizing the sections, in most cases, the same sections were reacted for CO to reveal the cortical laminae and V1 and V2 compartments, and then re-imaged under bright field illumination (1.25x-5x magnification). In 3 cases, axonal label location for 2-3 sections was determined by alignment to immediately adjacent CO-stained sections, rather than from the same section imaged for fluorescent label. In 4 cases, 1-2 sections were further immunoreacted to enhance the GFP signal strength, and then imaged a third time using both bright field and fluorescent channels to maintain a perfect alignment between the CO and label images.

### Quantification and Statistical Analysis

#### Analysis of Injection Site Location

Digitized fluorescent label and CO-staining from the same or adjacent sections were warped, using *IR-tweak* warping software (NCR toolset, Scientific Computing and Imaging Institute, University of Utah) by registering the radial blood vessels sequentially in a stack encompassing all cortical layers (for the tangentially sectioned cases). *IR-tweak* is an interactive, multithreaded, application for manual slice-to-slice registration. As control points are placed by the user in one image, their locations in the other image are estimated by the current thin-plate spline transform parameters ^64^. When the user corrects the locations of estimated points in the second image, the transform parameters are updated. CO-staining was used to assign injection sites to specific V2 CO stripes. To determine the CO stripe location of the injection sites, typically 3-6 images of serial CO sections were overlaid by aligning the radial blood vessels and merged in Adobe Photoshop. This approach reveals a more complete stripe pattern than any individual CO-stained section, in which the stripes can be incomplete. V2 injection sites were outlined through multiple images of serial injection sites, spanning the depth of the cortex; a composite injection site encompassing all outlines in individual sections was overlaid onto the merged CO stack, and its size and stripe location were determined. The size of the injections was defined as the region at the injection site where labeled cell bodies could be discerned, as the viral vectors we used infect neuronal somata. Dark CO stripes were classified as thick (or thin) if they appeared thicker (or thinner) than the two adjacent dark stripes in CO staining and coincided with thick (or thin) stripes as defined in the optical imaging maps (see above *Optical Imaging*), for cases in which imaging was available. To align CO stained tissue and fluorescent images of the injection sites to the optical maps, we warped (using *IR-tweak* software) the most superficial tangential tissue sections containing the surface vessels (which run parallel to the cortical surface) to the image of the cortical surface vasculature obtained under green light (the latter in register with the functional maps obtained under red light). Each subsequent section was then aligned sequentially to the top sections by aligning the radial blood vessels (which run perpendicular to the cortical surface) (**Supplementary Fig. 1, Figs. 2A-C, 3**).

#### Laminar Distribution of FB Axons: Quantitative Analysis

For each injection case used for the laminar analysis (n=4), the V1 laminar distribution of labeled V2 FB axons was analyzed in 3 tissue sections (cut perpendicular to the cortical layers), of which one section was selected near the center of the FB labeled field, where FB terminations are densest, and 1 section was selected on each side of this center section, equally spaced across the full extent of the labeled field in L4B (as the FB labeled field in this layer had a smaller tangential extent than the labeled fields in other layers, and we wanted the ROI chosen for analysis to include all layers of FB termination). Sections were imaged for GFP and tdT fluorescent label, stained for CO (to reveal layers), then re-imaged for both CO staining and fluorescent label. The layer boundaries were traced manually using CO staining. For each section selected for quantitative analysis, a ROI (500 µm wide, the full length of the cortex) was centered on the area that contained the densest fluorescent label. The ROI was further binned from top of the cortex to the white matter border into 100 bins. Individual pixel fluorescent intensity values were measured within each bin, and averaged to obtain the label intensity per bin. The different bin intensities were then normalized to the maximum bin intensity in that ROI. The binning of the ROI provided a FB label intensity profile across the cortical depth. The normalized label intensity in each bin was finally averaged across the 3 section ROIs and, again, normalized to the max average intensity to produce a single V1 laminar profile of the label intensity for each injection case. Population laminar profiles for FB arising from the same V2 stripe type were obtained by averaging the laminar profiles across cases with injections in the same stripe type. The cortical layer boundaries derived from the average layer thickness across CO-stained sections were then superimposed to the label laminar profile (right panels in **Fig. 1E,F**).

To determine whether viral injections in different V2 stripe types produced different laminar profiles, we performed statistical comparisons on a layer-by-layer basis. Because layers can vary in thickness across different sections, and therefore ROIs, in this analysis the same layer could contain slightly different numbers of bins across ROIs, each having an average pixel intensity value. The *n* for a given layer dataset consisted of the average intensity value of each bin in that layer across all ROIs, pooled across all cases with an injection in the same V2 stripe type. For across stripe statistical comparison of FB label intensity for each layer, we used a nonparametric test for two independent samples (Mann-Whitney U test - using the Matlab’s *ranksum* function), due to the varying number of bins per layer and the non-normal distribution of the data.

For each stripe group, we also statistically compared the resulting FB label intensity across the sublayers of L4 (4A, 4B, 4C). The *n* for the sublayer dataset consisted of the average intensity value of each bin in that sublayer across all ROIs, pooled across all cases with an injection in the same V2 stripe. For statistical comparison we used the Kruskal-Wallis test with a Bonferroni corrected α value of 0.017.

#### CO Compartment Distribution of FB Axons: Quantitative Analysis

This analysis was performed on tissue sectioned tangentially to the pial surface. For each injection case used for this analysis (n= 9 injections, 3 for each stripe type), digitized images of fluorescent label and CO staining were warped and aligned into a sequential stack through the depth of the cortex, as described above for the analysis of V2 injection site location. CO staining was used to identify layers, as well as the blobs and interblobs in the superficial layer sections. For each case, we selected for quantitative analysis at least one section (the one with the densest label) through each of the following layers: 1A, 1B, 2/3, 4B, 5/6. However, for some layers (particularly the thinner L1A, 1B and 4B), in some cases it was necessary to select for analysis two adjacent sections, because the cut was not perfectly tangential and therefore the label field in that layer was split between adjacent sections; this ensured that the entire labeled FB field for that layer was analyzed. To quantify the distribution of labeled FB axons across V1 CO compartments within each of the chosen sections, we extracted from a L2/3 CO-stained section in the aligned section stack the CO blob borders, blob centers and interblob center-lines (**Supplementary Fig. 3A**). Blob borders were extracted by processing digitized images of CO stained sections using low-pass and bell-shaped local histogram equalization filters (using Image Pro Inc. software); the latter filter gets rid of inequalities in CO staining intensity across the tissue by distributing the image histogram evenly around the center of the histogram intensity scale. This results in more uniform image intensity distribution across the imaged region (e.g. **Supplementary Fig. 2E**), from which one can, then, segment out the blobs as the darkest 33% of pixels across the gray scale image (yellow outlines in **Supplementary Figs. 2E, 3A**) ^22,65^. Blob centers were defined as the darkest spot within each blob (red dots in **Supplementary Fig. 3A**), as determined by the image analysis software; the latter were then visually inspected for accuracy. For blobs that showed significant elongation, we marked several centers; the latter generally coincided with multiple peaks in CO intensity along the length of the blob, having similar center-to-center spacing as the surrounding tissue (e.g. see several elongated blobs in **Supplementary Fig. 3A**). The centers of the interblobs were defined as the brightest pixels between adjacent blob centers (green outlines in **Supplementary Fig. 3A**).

For each injection case, we analyzed the CO map in V1 L2/3 encompassing the full tangential extent of the labeled FB fields across the layers. From this CO map we estimated a square region of interest (ROI; black box in **Supplementary Fig. 3A**) whose size encompassed the average blob and interblob size in the map. To estimate the ROI, for each blob in the CO map, we computed a mean blob center-to-interblob center (B_ctr_-I_ctr_) distance from radial distances measured at 20° increments around the blob center. A grand average B_ctr_-I_ctr_ distance was then computed from all relevant blobs’ mean B_ctr_-I_ctr_ distances. This grand average distance was doubled to obtain the ROI for that case. This ROI was then centered on each blob in the CO map, and the CO staining intensity was measured for each pixel within each blob ROI, summed across all blob ROIs, and normalized to the maximum summed intensity value, to produce a CO intensity heatmap for the case (**Supplementary Fig. 3B**). Mean blob diameter for the case was computed from all blobs used for the analysis and overlaid onto the CO heatmap (black circle in **Supplementary Fig. 3B**).

In a similar fashion we generated heatmaps of fluorescent FB label (GFP or tdT) intensity for each layer of FB termination, by measuring fluorescent intensity within the same size ROI used to compute the CO heatmap, centered on each blob in the CO map that overlaid the labeled FB field in that layer (**Supplementary Fig. 3C**). Therefore, the number of blobs used for this analysis could vary from layer to layer, depending on the extent and location of the labeled fields in each layer. Signal intensity was summed across all ROIs in the layer, and normalized to the maximum intensity, to produce a fluorescent intensity heatmap for that layer (**Supplementary Fig. 3D)**. A mean blob diameter specific to each layer was computed from all blobs used to generate each layer heatmap, and overlaid onto the layer heatmap (black circle in **Supplementary Fig. 3D**).

From the heat maps, we generated graphs of the normalized CO or fluorescence signal intensity as function of distance from the blob center (**Supplementary Fig. 3E**). Because the distribution of FB label was not homogenous all around the blobs (e.g. **Supplementary Fig. 3D**), these graphs were generated by measuring the average signal intensity in each of 20 bins spanning from the center of the blob (zero on the x axis) to the center of the interblob (1.0 on the x axis) within a 200 µm-wide window centered in the region of highest signal intensity in the heat map (black boxes in **Supplementary Fig. 3B,D**); signal intensity was finally normalized to the maximum bin intensity value. The region of highest intensity on the heat map was determined by measuring the average signal intensity within a 200µm-wide window rotated every 20° around the blob center, and selecting the window with the highest average signal intensity. As the latter was measured independently on the CO and fluorescence heatmaps, their respective windows of highest signal intensity did not necessarily match in location around the average blob center (e.g. in **Supplementary Fig. 3D**, the window is located at 90°, while in **Supplementary Fig. 3B** it is located at 290°).

To produce population heatmaps for each cortical layer (**Fig. 7**), the layer heatmaps from each case were scaled up to the largest heatmap across the relevant cases, summed and normalized to the maximum summed value. The population graphs in **Fig. 7** were generated from the population heat maps as described above for the individual cases (**Supplementary Fig. 3E**).

We statistically compared the location of densest FB label produced by injections in different stripe type. To this purpose, for each case layer heatmap, within the 200µm window of densest label, out of the 20 bins we selected for analysis those with >70% intensity. The locations of the selected bins relative to the blob center were compared layer by layer across stripe types. For statistical comparison across stripes we used a Kruskal-Wallis test with a Bonferroni corrected α value of 0.017. We also performed a statistical comparison by pooling all the selected bin locations across all layers for a given stripe type and comparing across stripe types, again using Kruskal-Wallis test with Bonferroni correction.

## Data and Code Availability

The Matlab code used to generate the heatmaps and perform data analyses in this study are found on GitHub at https://github.com/angeluccilab/streamspecificfeedback. The data that support the findings of this study are available from the corresponding author upon reasonable request.

## ACKNOWLEDGMENTS

We thank Kesi Sainsbury for expert technical assistance, and Dr. Jeff Yarch for help with some experiments. We also thank Drs. Stewart Shipp and Karl J. Friston (UCL) for helpful discussions on predictive coding and the role of feedback. Supported by grants from NIH (R01 EY026812, R01 EY019743, BRAIN U01 NS099702), NSF (IOS 1755431, EAGER 1649923), University of Utah Research Foundation and Neuroscience Initiative, to A.A., and from Research to Prevent Blindness, Inc. and a core grant from NIH (EY014800) to the Department of Ophthalmology, University of Utah.

## AUTHOR CONTRIBUTIONS

Conceptualization: all authors. Methodology: all authors. Software: S.T, M.S.H. Validation: S.T., F.F., A.A. Formal Analysis: S.T., M.S.H. Investigation: F.F., S.M., A.A. Writing-Original Draft: F.F., S.T., A.A. Writing–Review/Editing: all authors. Visualization: F.F., S.T. Supervision & Funding Acquisition: A.A.

## COMPETING INTERESTS

The authors declare no competing interests.

## FIGURE LEGENDS

**Supplementary Figure 1.**
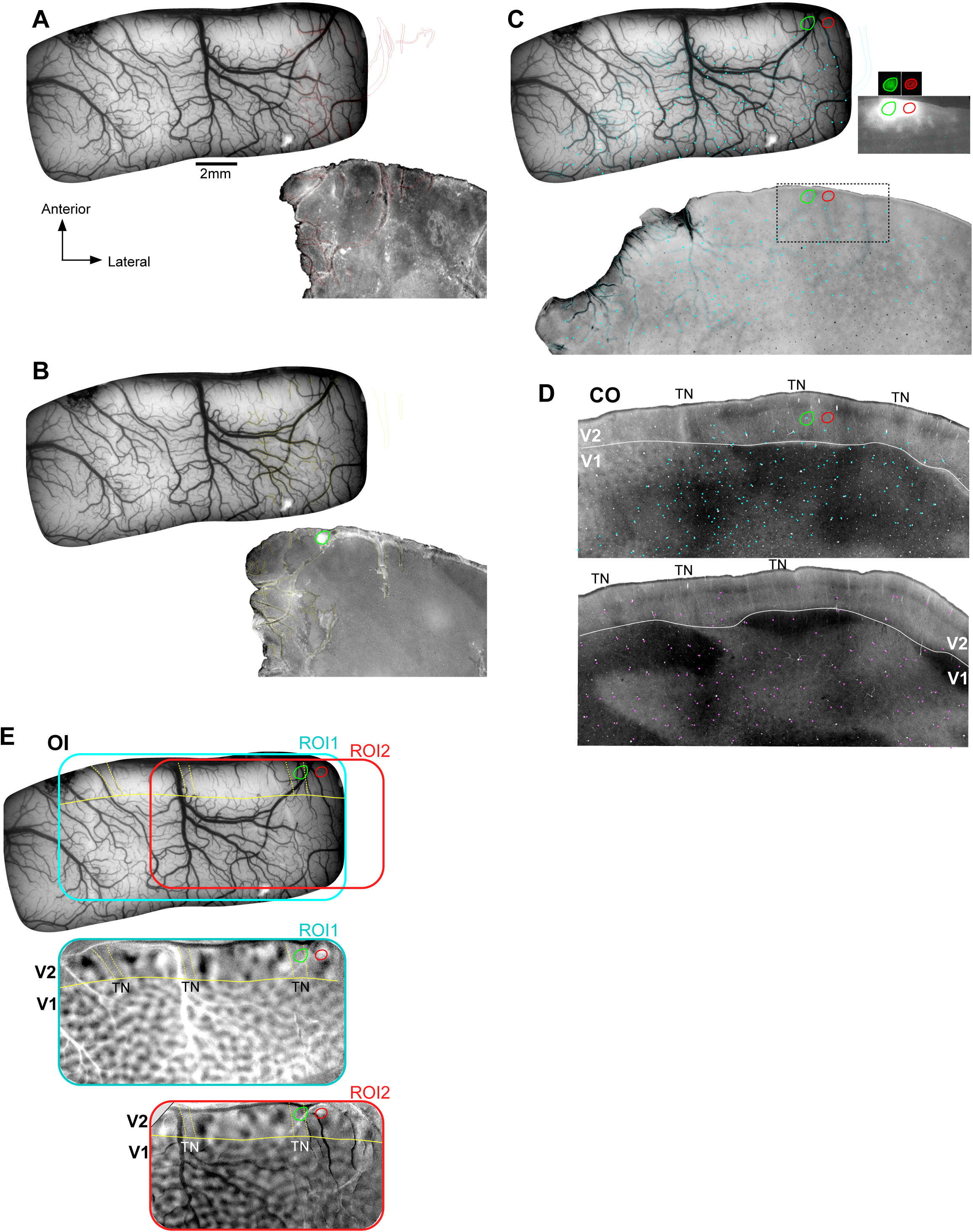
Alignment of histological sections to *in vivo* optical imaging (OI). **(A-C)** TOP: *In vivo* image of the cortical surface vasculature taken under green light illumination. BOTTOM: microscopic images of 3 superficial tissue sections (A most superficial and C deepest) containing surface, tangentially-running, vasculature. The superficial blood vessels (BVs) in the histological sections were warped to the BVs in the optical image. To demonstrate the alignment procedure, BVs in each tissue sections are outlined and shown overlaid to the *in vivo* optical image. *Cyan dots in (C)* mark the position of the radial BVs in the histological section, and are shown overlaid to the *in vivo* image, where they correspond to points where the surface BVs penetrate the cortex. *Inset in (C):* fluorescent microscope images of the viral injection sites from the same tissue section in the region inside the *black box*. The grayscale image was overexposed to visualize the radial BVs (used for alignment), which are marked as *cyan ovals*. The color images above it show the injection sites under correct light exposure (outlined), also shown at higher power in **Fig. 2C**. The injection outlines are shown superimposed to the optical image (TOP), the CO section in (D), and the orientation maps (E). **(D)** Two deeper tissue sections stained for CO to reveal the V2 stripes. The top section was warped to the section in (C) by aligning the radial BVs across serial sections (*cyan dots* mark the vessels from C); the bottom sectio n (240µm deeper) was warped to the top section whose radial BVs are marked as *pink dots. Dashed contours:* outlines of V2 stripes; *solid contour:* V1/V2 border. Due the sectioning plane being not perfectly tangential, the CO stripes are best visible in the lateral part of the top section and in the medial part of the bottom section. **(E)** Orientation difference maps from two different ROIs corresponding to the location of the *cyan and red boxes* at the top. *Yellow contours* outline the *TN* stripes and the V1/V2 border as revealed in the orientation maps.

**Supplementary Figure 2.**
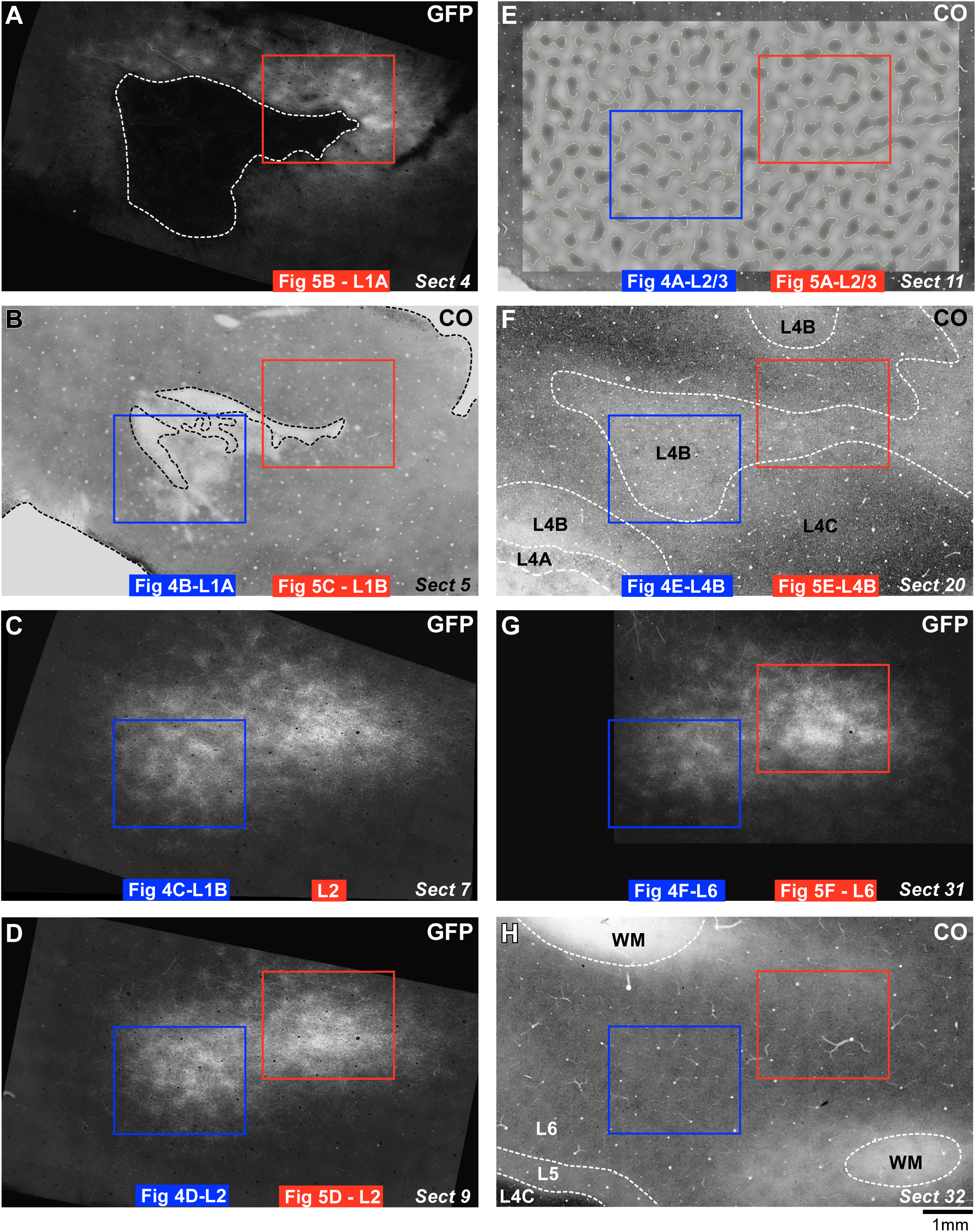
Case MK356RH. Low power view, and laminar locations, of the two terminal fields of FB connections arising from the thick (Left field) and thin (Right field) stripe injections shown in **Fig. 3**. *Blue* and *red boxes:* locations of the terminal FB field regions analyzed for the thick and thin stripe injection, respectively, and corresponding to the images shown in **Figs. 4-5**, as indicated below each respective box. To demonstrate the laminar location of the labeled terminal FB fields, we show CO-staining of the same section used for the GFP-label analysis (B) or of immediately adjacent sections (E,F,H). Sections in (A-H) are aligned and presented in sequence from the most superficial section (A), to the deepest (section # indicated at the bottom right corner in each panel). The sections in (F) is immediately deeper to the section in **Figs. 4E,5E**.

**Supplementary Figure 3.**
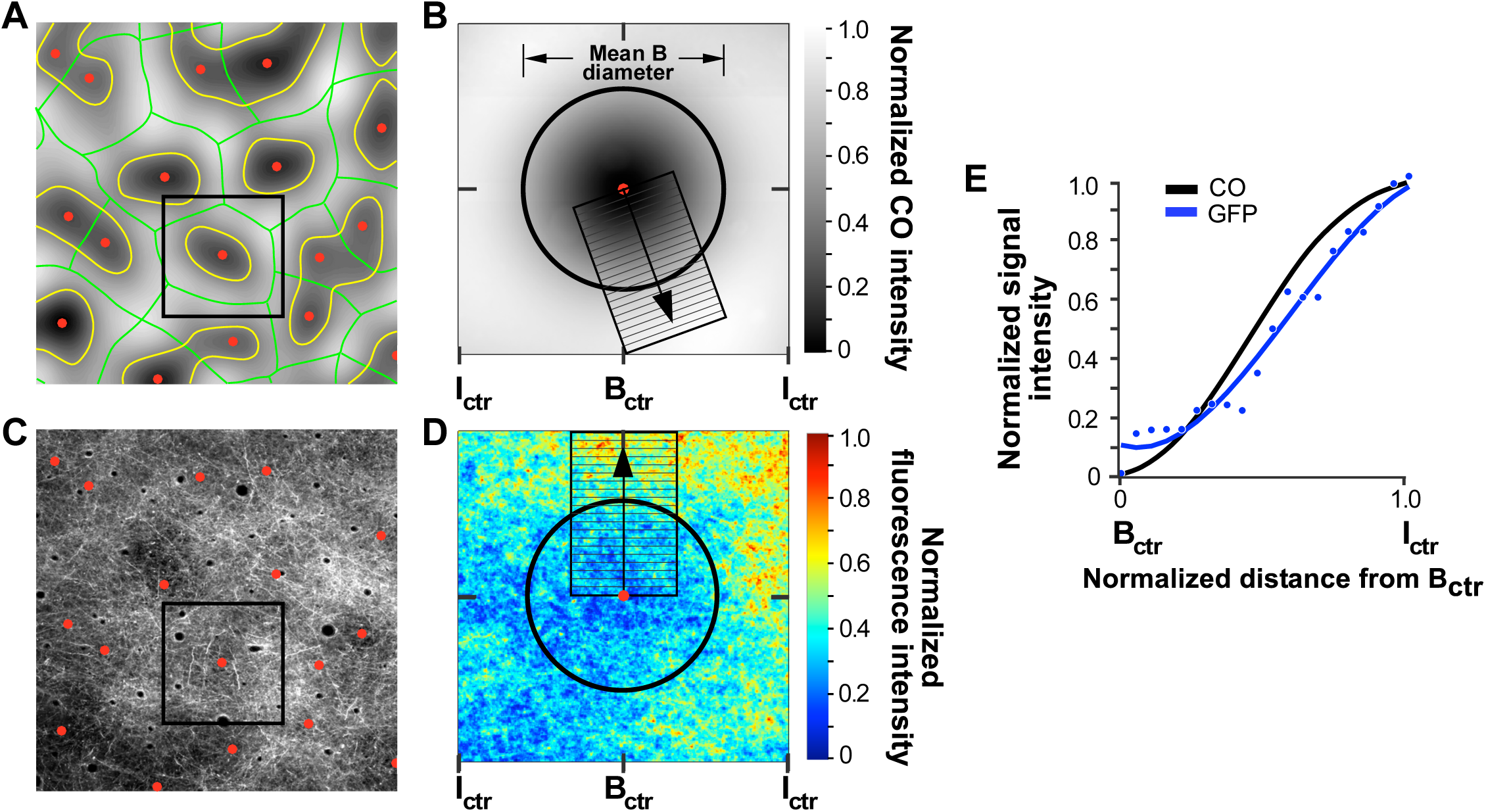
Quantitative analysis of the distribution of FB axons across V1 CO compartments. **(A)** Filtered CO stained section through L2/3 of V1. *Yellow and green contours and red dots* mark the blob borders, interblob and blob centers, respectively, extracted as described in the Methods. All blobs and interblobs in this map overlap the region containing labeled FB axons shown in (C) and were used to compute the heat map shown in (D). B*lack box:* an ROI, representing the average size of a blob and interblob in the larger CO map encompassing the full extent of the labeled FB fields across all layers. This ROI was centered on each blob in (A) and used to compute fluorescent signal intensity in (D). **(B)** Heat map of CO intensity, obtained by summing CO intensity across all blob ROIs for the case, and normalizing to maximum intensity. *Black circle*: mean blob diameter for all blobs used to compute the heat map. *Black box:* a 200 µm-wide window, centered on the region of highest CO intensity in the heat map, was divided into 20 bins from the blob center to the interblob center in the heat map. The mean signal intensity in each bin was normalized to the highest intensity value across all bins and plotted as a function of distance from the blob center (B_ctr_) in panel E (*black curve*). **(C)** Digitized image of GFP-labeled FB axons in a section through L2/3 of V1, vertically aligned to the CO section shown in (A). *Red dots* mark the blob centers from (A). The fluorescent signal intensity was measured in this section within the ROI from (A) centered on each blob overlaying the FB label in the layer. **(D)** Heat map of fluorescent signal intensity obtained by summing fluorescent intensity across all blob ROIs in (C), and normalizing to maximum intensity. *Black circle:* average blob diameter computed from all blobs in (A). The *black box* is the region of highest fluorescence intensity in the heat map, and was used to generate the plot of fluorescence intensity as a function of distance from the B_ctr_ in panel E (*blue curve*). **(E)** Plots of normalized CO and fluorescent signal intensities as a function of distance from the B_ctr_, measured within the black boxes shown in (B) and (D), respectively.

## Notes

https://github.com/angeluccilab/streamspecificfeedback

### Competing Interest Statement

The authors have declared no competing interest.

### Summary of Updates

Figures 2 and 3 revised; Supplementary Figures 2 and 3 added; contributing author added.

